# Scalable machine learning improves resistance prediction and identifies novel determinants in Mycobacterium tuberculosis

**DOI:** 10.64898/2026.04.25.720842

**Authors:** Mohammadali Serajian, Mahsa Lotfollahi, Oded Green, Kaleb Smith, Simone Marini, Mattia Prosperi, Christina Boucher

## Abstract

Multidrug-resistant and extensively drug-resistant *Mycobacterium tuberculosis* (MTB) represents a growing global health crisis, characterized by limited treatment options and high mortality rates. Rapid and accurate prediction of resistance profiles is critical to guide effective therapy and curb transmission. Whole-genome sequencing (WGS) offers promise for individualized resistance profiling, yet existing computational tools remain constrained by predefined mutation catalogs and prohibitive resource requirements for large-scale analyses. Here, we present AURA, a GPU-accelerated, pangenome-scale machine learning framework for de novo resistance prediction. Trained on 12,185 globally diverse MTB isolates, AURA predicts resistance to 13 first-line, second-line, and repurposed antibiotics with high precision and identifies 59 novel resistance-associated loci, including variants in *katG, pncA, rpoC*, and members of the *PE/PGRS* gene family. By enabling model training on an unprecedented genomic scale, AURA provides new insights into the genetic architecture of resistance and establishes a scalable platform for precision-guided therapy and global surveillance of MTB.

## Introduction

Tuberculosis (TB), caused by the *Mycobacterium tuberculosis* complex (MTBC), remains one of the leading infectious causes of morbidity and mortality worldwide [20]. Even though effective treatments have been available since the mid-20th century, TB continues to have a major impact, with 10.6 million new cases and 1.3 million deaths reported globally in 2022 [72, 74]. It is the world’s leading cause of death from a single infectious agent and among the top 10 causes of death [73]. Control efforts are challenged by persistent gaps in case detection, delays in diagnosis, inconsistent treatment adherence, and the accelerating emergence of drug resistance [17]. Multidrug-resistant TB (MDR-TB), denoted by resistance to rifampicin and isoniazid, accounts for approximately 410,000 new cases annually [70, 71]. Even more concerning is extensively drug-resistant TB (XDR-TB): MDR-TB with additional resistance to any fluoroquinolone and at least one Group A agent (bedaquiline and/or linezolid), which lowers treatment success to below 60%[8, 10, 61]. These trends undermine progress toward the World Health Organization (WHO) End TB Strategy. This strategy aims for a 90% reduction in TB incidence and a 95% decrease in mortality by 2035 [69].

The rise of *Mycobacterium tuberculosis* (MTB) drug resistance has underscored the urgent need for rapid and accurate antimicrobial resistance (AMR) profiling to guide effective therapy, support surveillance, and inform public health interventions [60]. However, antimicrobial susceptibility testing (AST) is still considered the gold standard, remains slow and resource-intensive, and requires several weeks due to the inherently slow growth of MTB in culture. Molecular diagnostics such as Xpert MTB/RIF and line probe assays (LPAs) have significantly reduced turnaround times. However, they are limited to the detection of predefined resistance-associated mutations within a narrow set of targets [44]. Although these assays are valuable in clinical practice, their scope does not extend to the identification of rare, lineage-specific, or novel mutations outside of their probe design.

Whole-genome sequencing (WGS) has emerged as a powerful tool in infectious disease genomics, allowing high-resolution insights into pathogen evolution, transmission dynamics, and AMR mechanisms. In the context of TB, WGS allows universal resistance profiling from a single assay, capturing the full genetic variation associated with drug resistance. Importantly, WGS can uncover previously unrecognized resistance determinants and provide information on the genomic epidemiology of MTB. Despite these advantages, integrating WGS into routine TB diagnostics faces significant hurdles, including laboratory infrastructure demands, data processing requirements, and the need for computational tools capable of analyzing large genomic datasets efficiently.

Several computational frameworks, such as TBProfiler [65], Mykrobe [27], and ResFinder [3, 19], have been developed to predict drug resistance from WGS data. These tools rely on curated mutation catalogs and alignment-based workflows, enabling rapid prediction for known resistance mutations. However, their dependence on predefined mutation panels limits their ability to detect resistance arising from novel genomic variants, structural rearrangements, or complex epistatic interactions. As drug pressure and global transmission contribute to the diversification of MTB genomes, there is a pressing need for computational approaches that can interrogate the pangenome comprehensively.

In this paper, we present AURA (Algorithmic Understanding of Resistance Associations), a GPU-accelerated de novo classifier designed for resistance prediction and feature discovery at the pangenome scale. Compared to CPU-based methods such as MTB++[54], AURA delivers more than 6,000-fold faster feature selection and 100-fold less memory footprint, enabling direct model training in the entire CRyPTIC cohort of 12,185 MTB genomes without reliance on alignment or predefined mutation panels. This capability supports not only rapid resistance prediction but also comprehensive biological discovery. More importantly, AURA emphasizes explainability by incorporating a feature analysis pipeline that aligns statistically significant genomic features to loci, facilitating biological interpretation and mechanistic insight.

Applying AURA to the CRyPTIC dataset uncovered 59 previously unreported resistance-associated loci. These findings validated known resistance genes such as *katG, pncA* and *rpoC*, while also highlighting genomic regions such as the *PE / PGRS* family, which are involved in immune evasion and host–pathogen interactions [13, 35, 53]. By enabling pangenome-scale machine learning (ML), AURA unifies predictive performance with biological interpretability. This capability is critical for advancing functional genomics studies aimed at validating resistance mechanisms and informing the development of next-generation diagnostics that detect a broader spectrum of resistance-conferring variants.

## Methods

### Dataset Description

All publicly available CRyPTIC [8, 61] MTB WGS and AST data were retrieved from the European Bioinformatics Institute (EBI) FTP repository. The WGS data comprise Illumina paired-end reads indexed by ERR accession numbers. The resulting dataset contained 12,185 isolates with paired genomic and phenotypic profiles. For each isolate, 13 antibiotics were tested and phenotypic drug resistance outcomes were reported. This dataset is available at the EBI^1^. The antibiotics include the first-line agents ethambutol, rifampicin, and isoniazid; the second-line agents amikacin, kanamycin, moxifloxacin, levofloxacin, ethionamide, and rifabutin; and the new or repurposed drugs (NRDs) linezolid, delamanid, clofazimine, and bedaquiline. Notably, “intermediate” phenotypes were reported only for ethionamide and ethambutol; in these cases, intermediate isolates were grouped with the resistant class to avoid misclassifying reduced-susceptibility strains as susceptible, which would bias the model toward overly optimistic predictions. Isolates lacking phenotypic susceptibility data for a given antibiotic were excluded from both model training and performance evaluation for that antibiotic-specific resistance outcome.

### Genome Assembly and *k*-mer Counting

Next, we ran Gerbil [16] on all 12,185 assembled genomes. Gerbil is a GPU-accelerated *k*-mer extraction tool. It uses a two-phase approach: the first phase reduces redundancy by grouping *k*-mers into super-mers using minimizers [50], while the second phase processes temporary files to count *k*-mer frequencies using hybrid CPU-GPU threading. Gerbil dynamically distributes workloads between CPU and GPU processors, ensuring efficient memory use and high throughput. Moreover, the software can output two main formats: binary and FASTA. As part of this GPU acceleration study, Gerbil was further modified to output in a CSV-format data frame, with one column for *k*-mers and another for their counts. This is to achieve compatibility and efficiency using the RAPIDS AI [49] framework. Furthermore, an additional filter was added to Gerbil, which excludes *k*-mers that exceed a threshold.

We processed the full genome set with Gerbil to extract all unique *k*-mers, which later form the columns of our feature matrix. During this step, we set the minimum k-mer count to 5 as the lowest practical value, since using 4 or 3 introduced many extremely rare k-mers that sharply expanded the feature space and caused GPU memory overflow, and we set the maximum to 6,500 as slightly more than half of the cohort to remove near-ubiquitous k-mers, noting that both thresholds are dataset-size dependent and should be adjusted for substantially smaller or larger datasets.

### Feature Matrix Construction

The feature matrix represents the frequency of each specific *k*-mer across a collection of genomes, providing a comprehensive view of *k*-mer distribution patterns. Here, the columns of the matrix correspond to *k*-mers and the rows correspond to the genomes, and thus a value of *c* in column *i* and row *j* of the matrix implies that *i*-th *k*-mer occurs in genome *j c* times. This matrix is the input to the classifiers. The feature matrices corresponding to the CRyPTIC dataset [61] are extremely large and sparse, with more than 99% of the elements being zero. Thus, we use a compressed sparse row (CSR) [22] representation to store the feature matrices, which efficiently encodes matrices using three lists: `data`, containing all non-zero values in the matrix; `columns`, storing the column indices of these values; and `rows`, indicating the starting index of each row within the `data` list, i.e., `rows`[i] represents the starting index in `data` for nonzero elements of the *i*-th row. The interval [`rows`[*i*], `rows`[*i* + 1]) contains the non-zero elements indices for row *i*. If row *i* has no nonzero elements, then `rows`[i] = `rows`[i+1]. We illustrate a CSR through the following example. Here, we have a sparse matrix A.

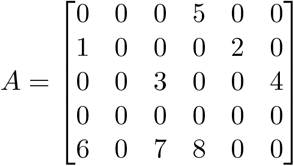

The corresponding CSR representation of A is as follows: `data` [5, 1, 2, 3, 4, 6, 7, 8], `columns` [3, 0, 4, 2, 5, 0, 2, 3] and `rows` [0, 1, 3, 5, 5, 8]. We briefly describe the entries of `rows`. The first entry, 0, appears because there are no non-zero entries before the first row. The second entry, 1, indicates that there is one nonzero entry (e.g., 5) in the first row. The third entry, 3, signifies that there are three non-zero entries (e.g., 5, 1, and 2) before the second row. Similarly, 5 appears next, indicating that there are five nonzero entries (e.g., 5, 1, 2, 3, and 4) before the third row. The next entry, 5, is repeated, meaning that the fourth row contains no nonzero entries. Finally, the last entry, 8, indicates that the eight non-zero entries are accounted for, with the fifth row contributing the last three entries (6, 7, and 8). This representation significantly reduces memory usage and computational overhead when handling large and sparse feature matrices. The CSR format stores only the non-zero elements along with their positions, enabling efficient row-wise access and matrix computations.

The key computational challenge lies in efficiently intersecting each genome-specific CSV file with the complete set of unique *k*-mers to generate the corresponding entries in the CSR representation—namely, the `data`, `columns`, and `rows` arrays for that genome. To accomplish this, we iterate through each of Gerbil’s 12,185 CSV outputs, intersecting each genome’s CSV file with the set of all unique *k*-mers using the cuDF merge function. Let *k*_1_, …, *k*_n_ be the counts of *k*-mers 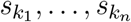 for the genome we are considering, and let 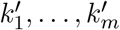 be the counts of *k*-mers 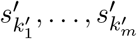 in the set of all genomes. This intersection transforms 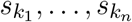 into entries in `columns`, `data`, and `rows`. To achieve this, we first create a hash map for 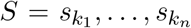. Next, we sort 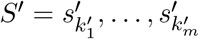 lexicographically and define the rank of each *k*-mer as its position in the sorted list, i.e., 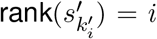 for all *i* = 1, …, *m*. We then query the hash map with each *s*^*′*^*k*^*′*^*i* to obtain the subset of *k*-mers, denoted as 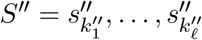, that are also present in *S*. We note that due to filtering, not every entry in *S*^*′*^ is guaranteed to be in *S*. We store two lists: the first contains the ranks of each 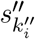, i.e., 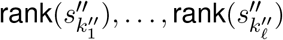, and the second contains their corresponding counts in *S*. These two lists are appended to the `columns` and `data` lists, respectively. Finally, the cumulative sum of existing elements is stored in the `rows` list to mark the row boundaries for each genome, ensuring efficient access to the feature matrix in CSR format. This structured representation allows for rapid querying and storage while preserving the sparsity inherent in the data set. The pseudocode for this construction is given in the Supplement.

We note that several design decisions were made in the creation of this algorithm that were made to increase the scalability of the algorithm. First, it may seem counterintuitive to create a hash table over the *k*-mers in the genome *S*^*′*^—rather than a hash table over all *k*-mers—but this substantially reduces the Video random-access memory (VRAM) usage of the algorithm. Next, we note that the CSR matrix lists (`data`, `columns`, and `rows`) are preallocated based on the estimated sparsity. If they reach capacity during construction, they are dynamically doubled, ensuring (1) insertions while maintaining efficient memory usage.

### Feature Selection

We conduct the univariate chi-square (*χ*^2^) test of independence in order to rank *k*-mers based on their *χ*^2^-scores and assess their relevance to resistance phenotypes, operating under the null hypothesis that the observed frequency of each *k*-mer is independent of the resistance phenotype— meaning, it should match the expected frequency if there is no association. By comparing the observed and expected frequencies, we identify *k*-mers whose distributions significantly deviate from the hypothesis of independence, suggesting their potential involvement in resistance. In this section, we describe these concepts and how they are implemented on GPU processors.

A contingency table is a matrix that summarizes the frequency distribution of *k*-mer frequencies across genomes relative to resistance phenotypes of the corresponding genomes, facilitating the analysis of their association. On CPU processors, Python libraries such as `SciPy` (`scipy.stats.chi2_contingency`) or `pandas` (`pandas.crosstab`) are used to compute the contingency table and assess statistical relationships. However, in large-scale genome-wide association studies, such as his study, the number of unique *k*-mers frequently reaches billions, necessitating the computation of contingency tables tens of billions of times. As a result, an iterative approach that processes each unique *k*-mer on CPU processors is computationally prohibitive, as identified in our previous study [54]. To address this limitation, we implemented the calculation of contingency tables using matrix multiplication on GPU processors.

To efficiently calculate the chi-square statistic for large datasets, we approximate the calculation by binarizing the CSR feature matrix **M**. This process can be done efficiently by binarization of `values` list of the CSR matrix. This transformation simplifies the problem by reducing the contingency table to a 2 × 2 matrix for each *k*-mer since the phenotypes have only two possible values. Using this approximation, the computation becomes highly efficient and scalable, allowing matrix multiplication-based calculations to replace iterative approaches. More formally, we let **L** denote the label or phenotype vector, where a value of 1 represents resistance, and a value of zero indicates susceptibility. The contingency table components (CT) is defined as follows:

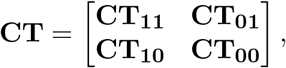

where **CT**_**11**_ denotes the number of samples with a resistant phenotype (**L** = 1) and a non-zero *k*-mers (**M** = 1); **CT**_**01**_ denotes the number of samples with a negative label (**L** = 0) and a non-zero feature (**M** = 1); **CT**_**10**_ denotes the number of samples with a positive label (**L** = 1) and a zero feature (**M** = 0); and **CT**_**00**_ denotes the number of samples with a negative label (**L** = 0) and a zero feature (**M** = 0). To compute these components efficiently, we use matrix multiplication to derive the contingency table values. Specifically,

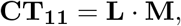

where · denotes matrix multiplication, **L** is the binary phenotype vector, and **M** is the binary feature matrix stored in CSR format.

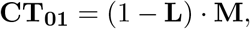

which computes the count of susceptible phenotypes with non-zero *k*-mers present.

For **CT**_**10**_, instead of explicitly calculating **L** · (1 − **M**), the computation can be optimized as:

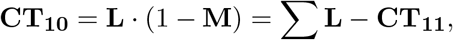

where ∑**L** represents the total count of genomes with resistant phenotypes.

Finally, **CT**_**00**_ is calculated as:

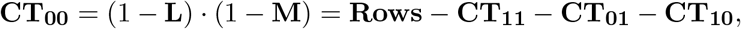

where **Rows** denotes the total number of rows in the feature matrix, which is equivalent to the number of all genomes.

This method avoids direct computation of dense matrices such as 1 − **M** and uses sparsity in **M**, significantly reducing both memory usage and computation time. By adopting this approximation and matrix multiplication approach, contingency table calculations become feasible for large-scale datasets.

After the calculation of the contingency table for all *k*-mers, the chi-square (*χ*^2^) scores are efficiently calculated on the GPU by leveraging matrix multiplication as follows. First, we calculate the marginal totals according to the following equations.

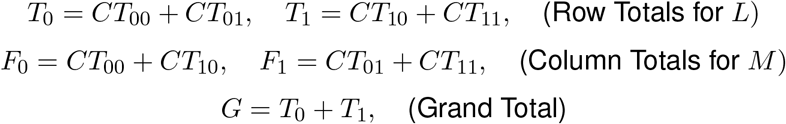

Next, the expected frequencies for each component are computed under the null hypothesis of independence as follows:

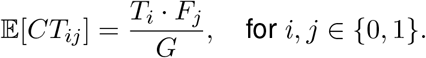

To account for the continuity of the chi-square test, Yates’ continuity correction is applied by adjusting the differences between observed and expected values as follows:

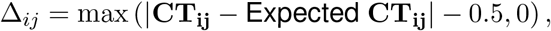

where *i, j* ∈ {0, 1}.

The chi-square score for each feature is computed as the sum of the normalized squared differences:

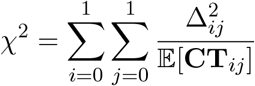

Lastly, to ensure numerical stability, a small constant *ϵ* is added to the expected frequencies to prevent division by zero.

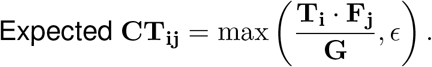

We parallelize the entire process on the GPU, leveraging matrix multiplication to efficiently compute all components of the contingency table and expected frequencies. This approach eliminates iterative loops and enables the simultaneous computation of chi-square scores for billions of features.

### Model Selection

A variety of ML algorithms have been applied to train AURA models, each offering distinct advantages that justify their inclusion. First, least absolute shrinkage and selection operator (LASSO) logistic regression (LR) uses an L1 penalty to manage a high-dimensional feature matrix, reducing irrelevant coefficients to zero and simplifying the model for better interpretation. Second, Random Forests build multiple decision trees to understand complex interactions in *k*-mer data, preventing overfitting and offering insights into the importance of features. Third, Naïve Bayes (NB) relies on conditional independence assumptions for efficient computation, making it effective for large, sparse *k*-mer datasets. Fourth, eXtreme Gradient Boosting (XGBoost) improves predictive accuracy in complex genomic tasks through optimized gradient boosting, parallelization, and advanced regularization. We note that the cuML library does not yet support CSR representation for training of some models, such as the LASSO LR, and random forest (RF) models; as a result, we had to use the dense format. However, this does not pose a significant issue, as feature selection drastically reduces the size of the feature matrix for each antibiotic drug.

### Performance Measures

We randomly partitioned the dataset of 12,185 MTB genomes into two independent subsets: 80% training and 20% testing. Each model was trained in the designated training set, followed by an evaluation in the independent test set. AURA predictive performance, in comparison with competing methods, was evaluated using F-1 score, recall, and precision on the test set. The competing methods are MTB++ [54], TBProfiler [45], ResFinder [3], KvarQ [58], and Mykrobe [27]. Lastly, we note that 6,224 of the 12,185 genomes were specifically used for the development of MTB++ [54] and were only included in the training set of AURA to ensure a fair comparison.

### Feature Analysis

AURA trained models contain a list of statistically significant features (*k*-mers) for each antibiotic that is used to predict antibiotic resistance. To assess the validity of these features, we aligned them to the MTB reference genome H37Rv (NCBI Reference Sequence: NC 000962.3) to identify loci associated with resistance to each resistance class. Alignments were performed using BWA [36], restricted to exact matches with the reference genome. The resulting BAM files were then intersected with the MTB GFF annotation file (NCBI RefSeq assembly: GCF 000195955.2) using BEDTools (version 2.30.0) [47]. For each antibiotic, numerous MTB loci fall within the feature set; we therefore focus on a curated subset defined by model-specific importance scores. In XGBoost, importance corresponds to feature weight—the number of times a locus-linked feature appears in tree splits—whereas in Random Forest, it reflects the mean decrease in Gini impurity across nodes. For Logistic Regression and Linear support vector machine (SVM), importance equals the absolute value of the model coefficients. To contextualize these findings, we document all loci that overlap the CRyPTIC GWAS loci results, and in case there is a locus captured by the ten most informative features of each model, we report them as novel associations. The term “association” is used advisedly to indicate that the loci identified here are provisional candidates for resistance; determining causality will require focused mechanistic studies. We report them now so they can be prioritized for further investigation in future work.

## Results

### Dataset Description

MTB sequence reads for all 12,185 isolates were assembled using SPAdes v3.15.3 [46]. Across isolates, the mean number of reads per genome was 4,305,537 (SD = 2,325,833). Assembly quality was assessed using QUAST [23]: the mean total assembly length (sum of contig lengths) was 4,488,463 bp (SD = 589,853 bp), the mean contig count was 645 (SD = 1,537), and the mean N50 was 97,027 bp (SD = 45,038 bp). The number of ambiguous bases (Ns) averaged 1,908 (SD = 1,056), with a mean NG50 of 97,941 bp (SD = 44,982 bp).

### The AURA Machine Learning Model

AURA is a GPU-accelerated ML framework developed using NVIDIA RAPIDS AI, an open-source suite of data science libraries for GPUs. AURA leverages NB, LASSO LR, SVM, RF, and XG-Boost to predict MTB drug resistance using WGS data. AURA is applied to CRyPTIC [61] data, which comprises WGS data of 12,185 human-host MTB isolates paired with AST for 13 antibiotics, including bedaquiline, clofazimine, linezolid, delamanid, amikacin, kanamycin, moxifloxacin, levofloxacin, ethionamide, ethambutol, isoniazid, rifampicin, and rifabutin. In order to evaluate the predictive performance of AURA, isolates were randomly stratified into 80% training and 20% testing for each antibiotic. AURA uses various sizes of *k*-mers extracted from assembled genomes, which are used as features. The extracted values of *k* are {9, 13, 17, 19, 25, 28, 31, 47, 61}. These *k*-mers are independently ranked for each antibiotic using the *χ*^2^ score calculated on the training data, and the first (2^21^) statistically significant features are used to train the predictive models.

All F1-scores in Table 1 were obtained using a per-antibiotic model-selection procedure in which k-mer length was optimized (not fixed across drugs). For each antibiotic, we split the data into 70% training, 10% validation, and 20% held-out test. We tuned k-mer length, the number of top-ranked k-mers retained (from an exponential series), and the classifier family (LR, RF, SVM, XGB, NB) using only the 70%/10% split, then selected the single configuration that maximized validation performance. Importantly, after selection, we did not change the split: we simply merged the same 70% training and 10% validation partitions (i.e., the exact samples already used for tuning) to retrain a final model with the chosen settings, and then evaluated it once on the untouched 20% held-out test set, and performance is measured.

**Table 1:** Comparison of F1-scores (%) for AURA and competing methods. Some baseline tools do not cover every drug examined in this study; these missing values are marked “N/A.” “Balance” reflects the proportion of resistant isolates, expressed as a percentage.

|  | Resistance Class | Balance | AURA | ResFinder | TBProfiler | Mykrobe | KvarQ | MTB++ |
| --- | --- | --- | --- | --- | --- | --- | --- | --- |
| First | Isoniazid | 48.48 | 94.69 | 94.37 | **95.34** | **95.34** | 93.41 | 94.22 |
|  | Rifampicin | 38.43 | 90.80 | 91.54 | **92.21** | 91.74 | 89.77 | 89.42 |
|  | Ethambutol | 31.38 | 81.00 | 81.64 | **81.89** | 81.85 | 77.12 | 77.88 |
| Second | Rifabutin | 36.74 | **91.19** | N/A | N/A | N/A | N/A | 88.86 |
|  | Ethionamide | 22.35 | **76.14** | 68.34 | 72.60 | 68.34 | N/A | 68.11 |
|  | Levofloxacin | 17.37 | 84.22 | N/A | **84.82** | 84.69 | N/A | 84.38 |
|  | Moxifloxacin | 14.28 | 74.12 | N/A | 73.83 | **75.00** | N/A | 70.23 |
|  | Kanamycin | 9.29 | **73.18** | 71.82 | 71.72 | 72.77 | N/A | 72.24 |
|  | Amikacin | 7.27 | 81.23 | **81.48** | 80.25 | 79.38 | N/A | 79.00 |
| NRDs | Clofazimine | 4.27 | **18.58** | 0.00 | 0.00 | N/A | N/A | 2.63 |
|  | Delamanid | 1.53 | **13.79** | N/A | 10.53 | 10.53 | N/A | 0.00 |
|  | Linezolid | 1.26 | **43.90** | **43.90** | 42.86 | 40.00 | N/A | 22.86 |
|  | Bedaquiline | 0.89 | **23.08** | 0.00 | 7.69 | N/A | N/A | 0.00 |

Lastly, for each antibiotic, AURA is compared against competing tools—including TBProfiler [45], ResFinder [19], Mykrobe [27], KvarQ [58], and MTB++ [54]—by executing all methods on the same test set. Table 1 reports F1-score comparisons across models. Precision–recall and specificity–sensitivity metrics for AURA and other methods are provided in Tables 5 and 6, respectively.

### AURA enables robust resistance profiling and advances comprehensive prediction for MDR/XDR-TB

Across the 13 antibiotics, AURA outperformed or closely followed the existing state-of-the-art tools, achieving the highest F1 scores for most second-line and NRDs, particularly rifabutin, ethionamide, kanamycin, linezolid, delamanid, clofazimine, and bedaquiline, which are critical for the treatment of XDR-TB and MDR-TB. Accurate prediction of resistance for these drugs is essential to guide effective therapeutic decisions. See Table 1, Figure 3.

**Figure 1:**
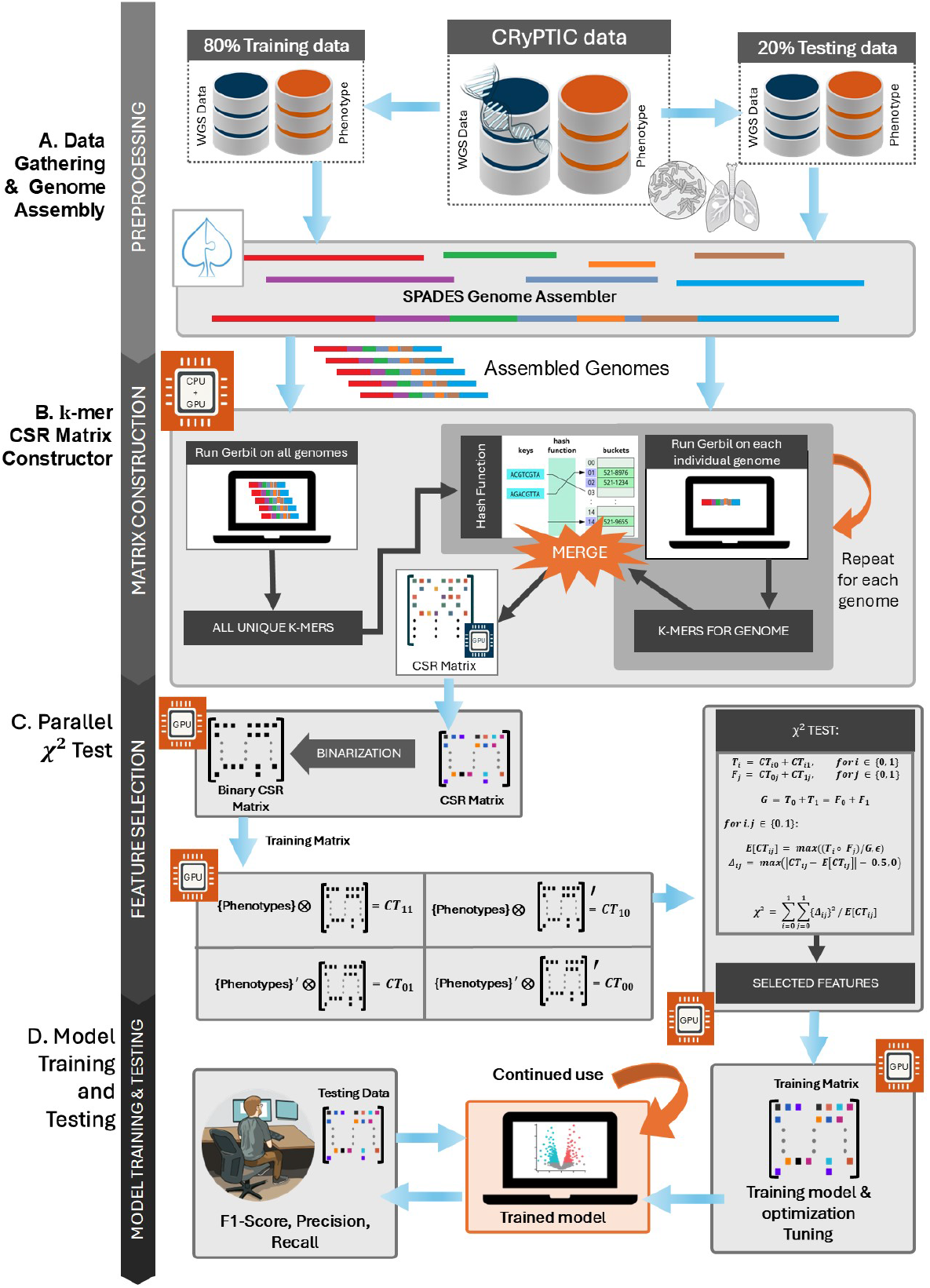
Overview of the AURA pipeline for resistance prediction in MTB. (A) Preprocessing: WGS data and the corresponding drug-resistance phenotypes from 12,185 MTB isolates [61] were divided into training (80%) and independent test (20%) datasets. The isolates were assembled *de novo* using SPAdes to generate contigs, ensuring comprehensive genomic representation for downstream resistance analysis. (B) Matrix construction: KMX enumerated *k*-mers (*k* = 9–61) across genomes using GERBIL, filtered them by frequency to reduce noise, and encoded per-genome profiles as compressed sparse row (CSR) matrices for efficient high-throughput analysis. (C) Feature selection: GPU-accelerated *χ*2 univariate test of independence identified the *k*-mers most strongly associated with drug-resistance phenotypes in the training data. The most statistically significant features, which are predictive of clinically relevant drug resistance, will be fed to ML models. (D) Model training and evaluation: Selected features were used to train ML classifiers (XGBoost, SVM, RF, LR, NB). For each antibiotic, optimized models were validated on the independent test dataset and the resulting F1-score, precision, and recall to assess predictive performance for potential clinical application.

**Figure 2:**
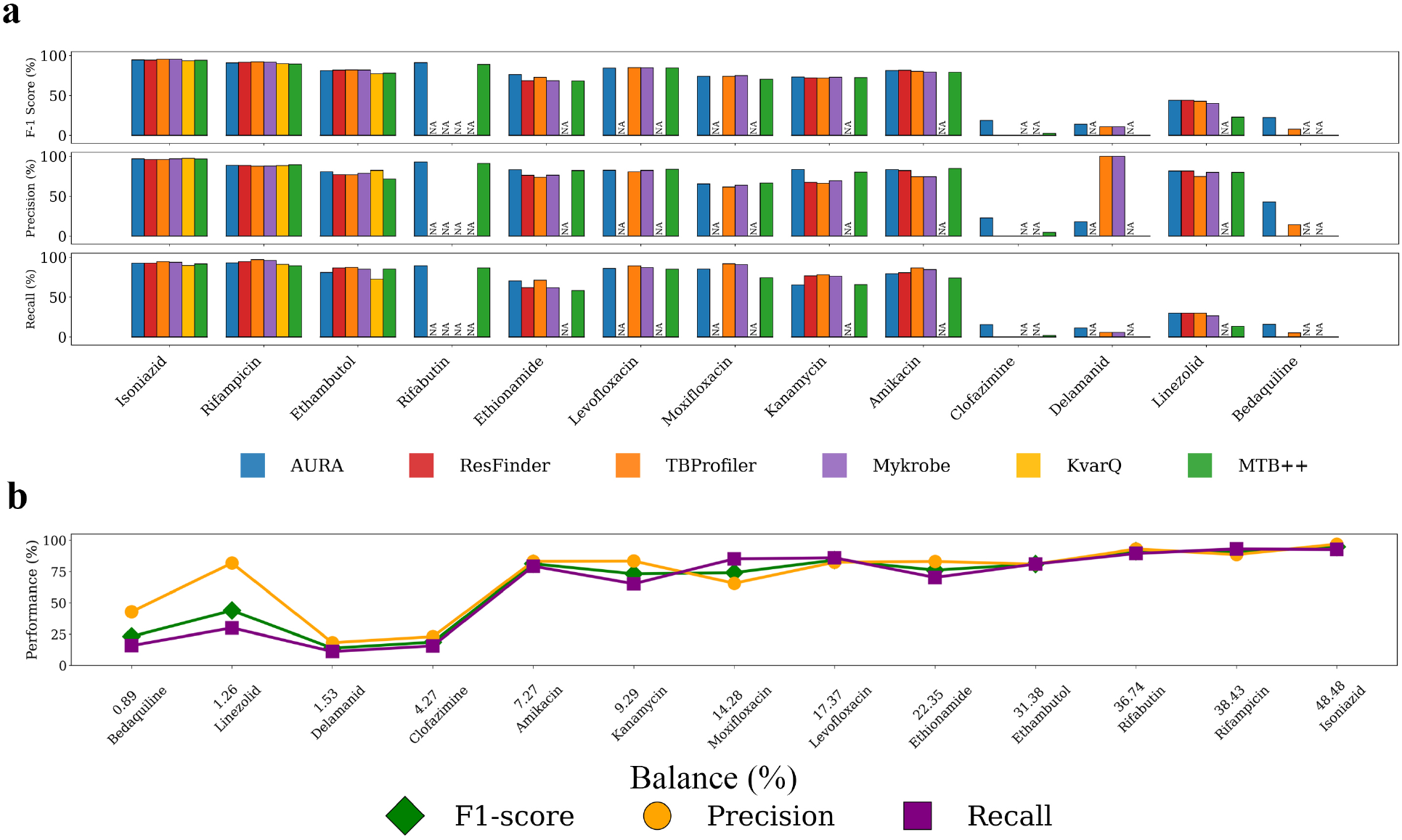
**a**, Per-antibiotic precision, recall, and F1-score for first-line, second-line, and NRDs achieved by AURA’s final optimized models and five reference tools (TBProfiler, ResFinder, KvarQ, Mykrobe, MTB++). This optimization is over both *k*-mer length and the ML models. **b**, Effect of class balance on predictive performance across antibiotics. For each antibiotic, the plot displays precision, recall, and F1-score achieved by the model for that specific drug. The percentage below every antibiotic gives the class balance—the proportion of resistant isolates among all isolates for that antibiotic.

**Figure 3:**
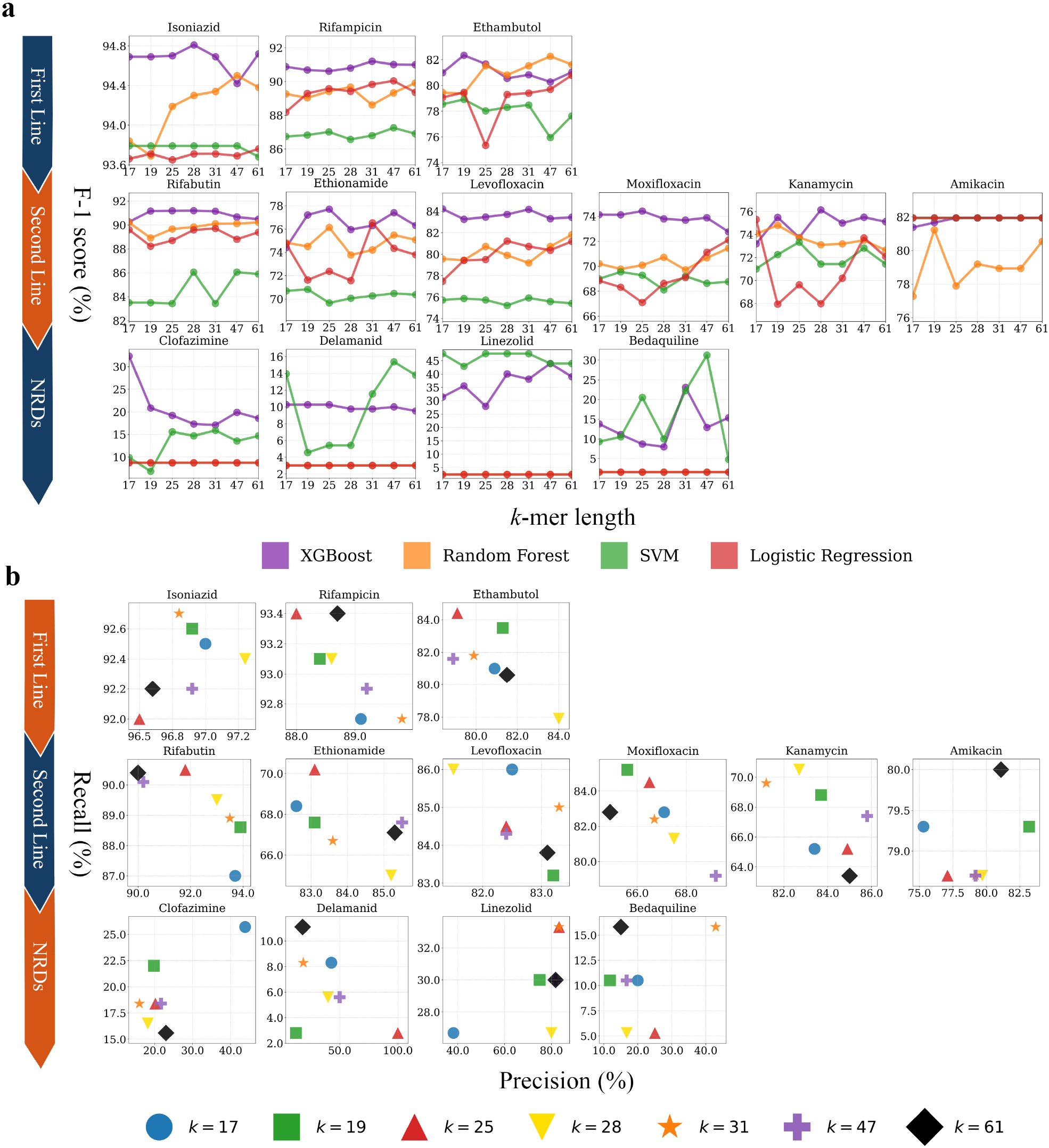
**a**, F1-score trajectories of AURA across six *k*-mer lengths (*k* ∈ 17, 19, 28, 31, 47, 61) for first-line, second-line, and NRDs. Each curve shows the performance of the selected classifier (XGBoost, SVM, RF, or LASSO LR) for an antibiotic. Results are from the held-out test set; *k* and model class were not optimized, while other hyperparameters were tuned on the validation set. **b**, Scatter plot of test-set precision–recall pairs for six *k*-mer lengths. At each *k*, the point corresponds to the classifier class (chosen from XGBoost, SVM, RF, and LASSO LR) selected on the validation set, together with its tuned hyperparameters. The model class and remaining hyperparameters were optimized on the validation set, while *k* was not optimized. Precision and recall are reported on the held-out test set.

A key factor contributing to this improved performance is the computational efficiency of AURA, which allowed extensive optimization of the model on different *k*-mer lengths for each antibiotic. Our analysis revealed a substantial variation in predictive performance depending on the *k*-mer length. Interestingly, several antibiotics reached peak F-1 score at shorter lengths of *k*-mer. For example, ethambutol (F1 = 81.00%), levofloxacin (F1 = 84.22%), and kanamycin (F1 = 73.18%) performed best at *k* = 17. Slightly longer *k*-mers yielded optimal results for moxifloxacin (F1 = 74.12% at *k* = 19), amikacin (F1 = 81.23% at *k* = 19), ethionamide (F1 = 76.14% at *k* = 25), rifabutin(F1 = 91.19% at *k* = 28), rifampicin(F1 = 90.80% at *k* = 28). Moderate *k*-mer lengths, particularly *k* = 31, proved ideal for drugs such as isoniazid (F1 = 94.69%), and bedaquiline (F1 = 23.08%). Even longer *k*-mers were optimal for clofazimine (F1 = 18.58%, *k*=61) and delamanid (F1 = 13.79% at *k* = 61) and linezolid (43.90% at *k*=47).

Across all antibiotics, we observed a consistent trend: the predictive precision improved substantially from *k* = 13. Performance at very short *k*-mer lengths (e.g., *k* = 9) remained low, likely due to insufficient sequence complexity and lack of discriminative genomic signals. Once *k* exceeded this threshold, the performance of the model increased dramatically, indicating that *k* = 13 represents a critical inflection point for the emergence of informative signals. For most first-line and second-line antibiotics, optimal performance was typically achieved within the range of *k* = 17 to *k* = 31, balancing predictive power with computational efficiency.

Even in resistance classes such as isoniazid, rifampicin, and ethambutol, where AURA did not produce the absolute highest score, it remained within 1–2% of the top performing tool, highlighting its consistently competitive performance. Furthermore, among the methods evaluated, only AURA and MTB++ produced resistance predictions for the 13 antibiotics profiled by the CRyPTIC consortium [61]. Competing tools often lacked modules for newer or second-line drugs, which therefore appear as N/A in Table 1. This comprehensive coverage demonstrates the versatility of our approach and its practical value. Although some tools, such as TBProfiler and ResFinder, can analyze some additional antibiotics, resistance classes fall outside the scope of AST in the CRyPTIC study [9, 10]; therefore, no models are developed for them.

Figure 3 further demonstrates the robustness of AURA across *k*-mer values. XGBoost and SVM classifiers delivered consistently high performance with minimal variation across the tested conditions. A key advantage of AURA is its ability to tailor the *k*-mer length to each antibiotic, striking an effective balance between overall model generalization and drug-specific refinement.

In summary, these findings position AURA as a highly accurate, adaptable, and comprehensive resistance prediction framework. Its ability to support a wide range of antibiotics, optimize predictions by tuning *k*-mer, and maintain high accuracy makes it a powerful tool for clinical and epidemiological applications.

### AURA expands resistance catalogs with novel genetic associations

To validate the predictive characteristics of AURA, we aligned statistically significant *k*-mers from trained models with the MTB reference genome using exact matching, identified resistance-associated loci via BEDTools [47], and prioritized them according to model-specific importance scores. This analysis recovered 151 resistance-associated loci previously reported by CRyPTIC [61], and identified 59 novel associations. See Table 2. We highlight a subset of the most biologically and clinically compelling associations identified by our analysis.

**Table 2:**
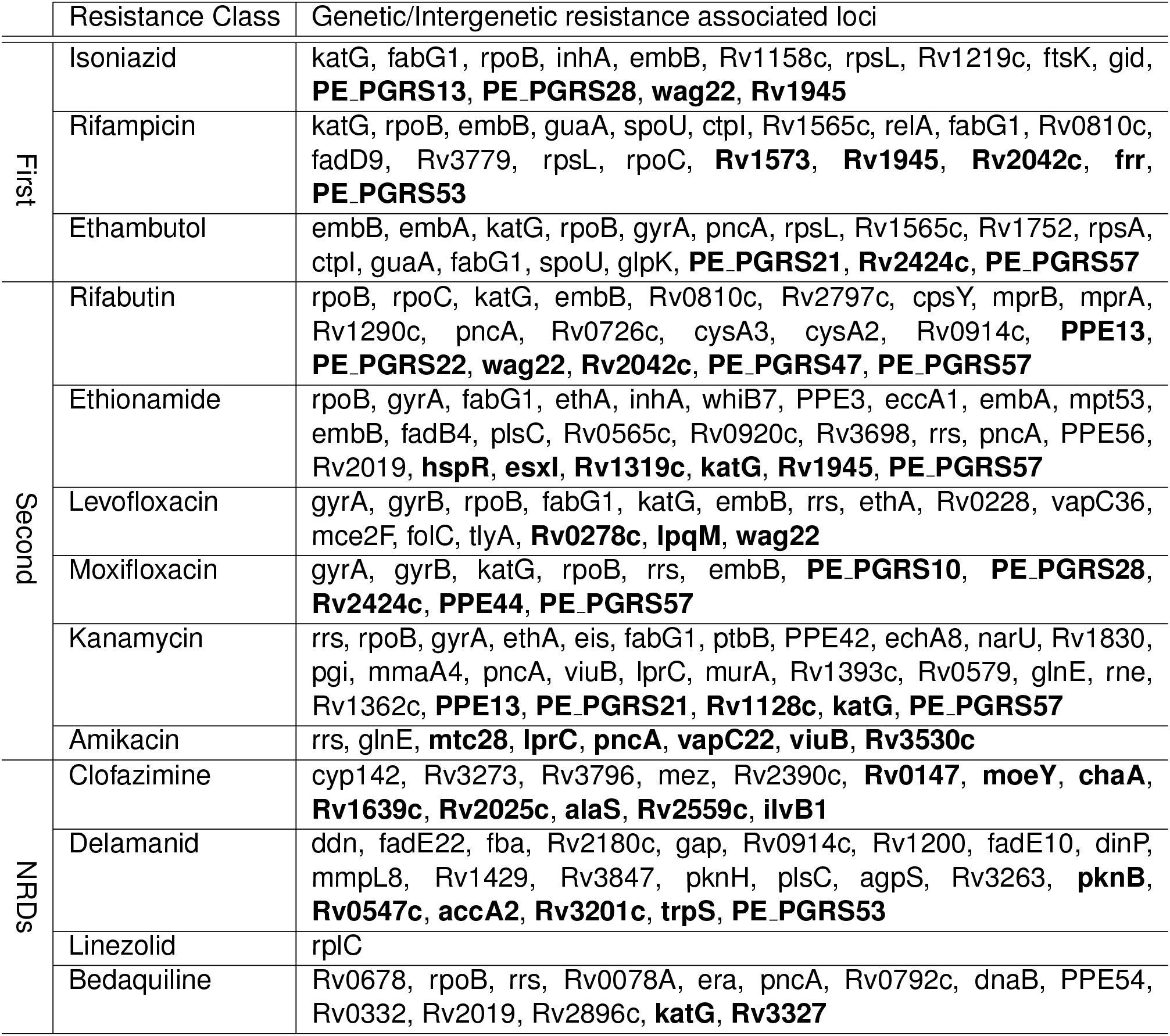
Genomic loci associated with antibiotic resistance identified by AURA, in comparison with those reported in the CRyPTIC GWAS loci results. Loci in **bold** represent associations identified by AURA that are not listed in the CRyPTIC GWAS loci results [61]; these may reflect co-resistance/co-treatment correlations, linkage, or population structure and should be interpreted as candidates for follow-up rather than established resistance determinants. Each resistance class includes both protein-coding genes and intergenic regions implicated in MTB drug resistance.

The *rpoB* locus emerged as the key determinant of resistance in MTB, showing associations with resistance to 9 of the 13 antibiotics tested—closely reflecting the patterns observed in the CRyPTIC GWAS loci results. This gene encodes the RNA polymerase *β*-subunit, which is essential for bacterial viability and pathogenicity [39, 40, 62].

As noted by Farhat et al. [18], recent clinical studies underscore the strong association between mutations in *rpoB* and resistance to rifampicin and rifabutin. In particular, mutations within the rifampicin resistance–determining region (RRDR) of *rpoB* (most notably S450L) have been shown to confer resistance to rifampicin. However, such mutations often come at a fitness cost as they can destabilize the RNA polymerase complex and affect transcriptional efficiency. Mutations in *rpoC* have been described as compensatory alterations that help restore the fitness cost associated with *rpoB*-mediated resistance to rifampicin, rather than acting as primary resistance determinants [14]. The *rpoC* locus encodes the *β*^*′*^-subunit that interfaces directly with the *β*-subunit. Variants at codons V483, L516, and N698 restore polymerase stability without adding further resistance and are enriched in multidrug-resistant (MDR) strains carrying S450L [14, 59, 67]. AURA feature analysis revealed *rpoC* to be exclusively associated with rifampicin and rifabutin resistance.

In addition to *rpoB* and *rpoC*, which mediate resistance to rifampicin and rifabutin through direct and compensatory mechanisms, our analysis identified *katG* as the second most frequently implicated locus across multiple antibiotics. In agreement with CRyPTIC, *katG* was associated with isoniazid, ethambutol, rifampicin, rifabutin, levofloxacin, and moxifloxacin. AURA further revealed *katG* novel associations with ethionamide, kanamycin, and bedaquiline. The *katG* gene encodes a bifunctional catalase–peroxidase that is indispensable for bioactivating the pro-drug isoniazid (INH). Once isoniazid enters the cytoplasm of MTB through passive diffusion [63], *katG* oxidizes it to form an isonicotinoyl radical, which then couples with NAD^+^ to generate the covalent INH–NADH adduct. This adduct binds tightly to the active site of the enoyl-ACP reductase *InhA*, thereby blocking the synthesis of mycolic acids, essential components of the mycobacterial cell wall, leading to cell death. Mutations in *katG* constitute the main genetic mechanism of isoniazid resistance, with Ser315Thr substitution being the most prevalent among clinical isolates [63].

The *pncA* gene emerged as a prominent and recurrent feature predictive of resistance to ethambutol, rifabutin, ethionamide, kanamycin, and bedaquiline, reinforcing prior associations reported by CRyPTIC. In addition, AURA identified an association between *pncA* and amikacin resistance. *pncA* is well established as the main genetic determinant of pyrazinamide resistance, and sequencing-based detection of *pncA* variants is often more reliable than phenotypic drug susceptibility tests, which may not capture low-level resistance conferred by dispersed nonsynonymous mutations across the gene [30, 41]. In vitro testing for pyrazinamide is technically challenging, costly, and time-consuming, as the drug is only active under acidic conditions, leading to false resistance rates as high as 70% [4, 5, 30].

The *pncA* encodes pyrazinamidase, a non-essential intracellular enzyme that converts the prodrug pyrazinamide into pyrazinoic acid, which is its active form[37, 52, 75]. This activation step supports the sterilization activity of pyrazinamide against persistent bacilli and MDR-TB [25, 30].

Mutations in *pncA* alter the folding, stability, or activity of pyrazinamidase, so pyrazinoic acid is not produced, and pyrazinamide becomes ineffective [56]. Consistent with this mechanism, most pyrazinamide-resistant clinical isolates harbor *pncA* mutations [29, 56]. Resistance-conferring mutations in *pncA* are scattered throughout the gene (unlike the defined hotspots in *rpoB* and *katG*), so comprehensive sequencing remains the most dependable method for detecting them [30, 37, 41]. Although alterations in *pncA* have not yet been shown to drive resistance to drugs other than pyrazinamide, statistical associations with several additional antibiotics have been reported [54, 55, 61], highlighting an area that deserves further investigation.

Moreover, we identified several genes belonging to the *PE PGRS* (proline–glutamic acid polymorphic GC-rich sequence) family as novel candidates associated with antibiotic resistance. The *PE PGRS* gene family constitutes a distinctive subgroup of *PE* proteins, which is conserved throughout MTB and selected other mycobacterial species [7, 35]. In our analysis, *PE PGRS57* (Rv3514) was notably prominent, showing novel associations with resistance to five antibiotics: ethambutol, rifabutin, ethionamide, moxifloxacin, and kanamycin. Supporting the importance of *PE PGRS57*, Seo et al. [53] recently identified this gene as the only locus consistently mutated in multiple strains resistant to a newly discovered class of small-molecule antimicrobials (PPs). Specifically, a recurrent Asp-to-Gly substitution within *PE PGRS57* conferred resistance, further implicating it in resistance mechanisms. Additional research has shown that other family members, such as *PE PGRS47* and *PE PGRS41*, contribute to immune evasion by suppressing host autophagy and antigen presentation [13, 51, 76]. Comparative genomic studies demonstrate that despite their antigenic variability, *PE PGRS* genes remain highly preserved and subject to purifying selection, underscoring their critical functional roles in MTB biology and disease pathogenesis [11, 43]. Collectively, these findings highlight the previously unrecognized involvement of the *PE PGRS* genes in antibiotic resistance and underscore their potential as candidates for further functional studies and therapeutic interventions in combating MDR-TB.

Lastly, it must be noted that the accuracy of *PE PPE* and *PE PGRS*–related association reported in this analysis is dependent on the quality of the SPAdes short-read assemblies in these highly repetitive, low-mappability regions, and any associated k-mers should therefore be interpreted cautiously.

### AURA leverages GPU acceleration for training on the largest TB resistance dataset to date

Previous computational frameworks for MTB resistance prediction have faced severe scalability barriers. Tools like MTB++ [54] rely on CPU-parallelized workflows, which, while effective on small cohorts, require prohibitive memory and storage resources when applied to larger genomic datasets. Notably, MTB++ demands over 2 TB of disk space and hundreds of gigabytes of random-access memory (RAM) to process even moderate datasets, limiting its utility for population-scale model development. In contrast, AURAeliminates these constraints and, to our knowledge, is the first GPU-accelerated framework to enable *de novo* training on the largest assembled tuberculosis WGS dataset that includes resistance phenotypes for NRDs. We benchmarked AURA against MTB++ across 3 key computational stages: feature matrix generation, feature selection, and classifier training, using identical datasets (Table 3).

**Table 3:** Comparison of AURA and MTB++ pipelines across three stages of resistance prediction for isoniazid (INH). Runtime is formatted as Days:Hours:Minutes:Seconds. A dash (–) indicates negligible usage or not applicable.

| Computational Stage | Method | Run-time | Main Mem (GB) | VRAM (GB) |
| --- | --- | --- | --- | --- |
| Feature Matrix Creation | MTB++ | 02:13:36:27 | 581.60 | – |
|  | AURA | 00:08:37:56 | 5.47 | 21.05 |
| $\chi^2$ Feature Selection | MTB++ | 02:19:39:40 | 56.48 | – |
|  | AURA | 00:00:00:39 | 13.53 | 29.76 |
| Classifier Training | MTB++ | 00:00:35:16 | 126.80 | – |
|  | AURA | 00:00:11:57 | 13.75 | 42.74 |

During feature matrix generation, MTB++ executed on 100 AMD EPYC 75F3 CPU cores required 2 days, 13 hours, 36 minutes (02:13:36:27), peaking at 581.60 GB RAM and generating 5.28 TB of intermediate artifacts. In contrast, AURA, running on a single AMD EPYC 75F3 core with an NVIDIA A100 GPU with 80 GB VRAM, completed this task in just 8 hours, 37 minutes (00:08:37:56), using 5.47 GB RAM, 21.05 GB VRAM, and only 13.4 GB disk space, representing a 6.2 times speedup and a 99.7% reduction in disk usage.

Feature selection, a critical bottleneck in prior workflows, was dramatically accelerated by AURA’s GPU implementation. MTB++ required nearly 3 days (02:19:39:40) and 56.48 GB RAM to process a single antibiotic (isoniazid), whereas AURA reduced this to 39 seconds (00:00:00:39) with 13.53 GB RAM and 29.76 GB VRAM, eliminating the need for disk I/O and achieving a 215× speedup.

In the classifier training stage, MTB++ required 35 minutes (00:00:35:16) and 126.80 GB RAM for the isoniazid model. AURA completed training in 11 minutes, 57 seconds (00:00:11:57) with 13.75 GB RAM and 42.74 GB VRAM, offering an approximate 3 times runtime reduction and over 9 times lower RAM usage.

Beyond training, AURA’s scalability extends to real-time resistance prediction across datasets of increasing size (1,000 to 10 million reads). While MTB++ showed modest scaling (3.9 seconds for 1,000 reads to 64.7 seconds for 10 million), AURA maintained competitive inference speeds, rising from 30.7 seconds to 185 seconds while individually optimizing *k*-mer lengths for each antibiotic. This performance outpaced all other tools tested, including ResFinder [3], Mykrobe [27], TBProfiler [45], and KvarQ [58], which exhibited steep runtime increases at scale (Figure 4). Importantly, all methods maintained memory footprints of less than 2 GB during inference.

**Figure 4:**
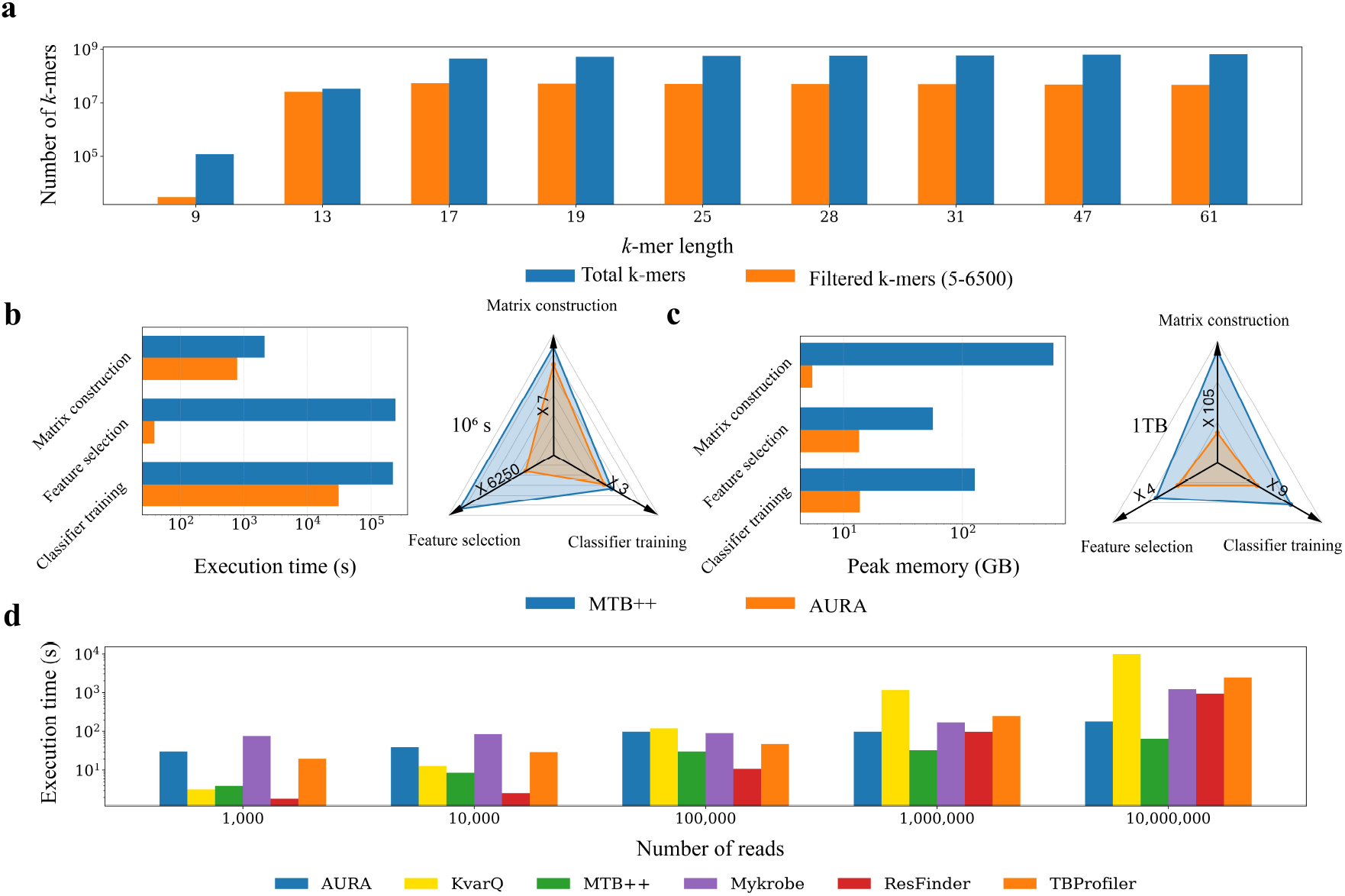
**a**, Expansion of the *k*-mer search space with increasing *k*. The blue curve reports the total number of *k*-mers identified in 12,185 MTB genomes, while the orange curve shows the subset retained after frequency filtering (occurrence ≥ 5 and ≤ 6,500). **b, c**, Log-scaled equilateral triangular polar plots comparing resource consumption for de novo training of MTB++ (blue) and AURA (orange). Panel B shows execution time for the three analysis stages—feature-matrix construction, feature selection, and classifier training, and Panel C shows similar measurements for peak RAM usage. **d**, depicts the execution time across increasing input sizes for all profilers. Input format was FASTQ for ResFinder [3], Mykrobe [27], KvarQ [58], and TBProfiler [45], and assembled FASTA for trained AURA and MTB++ [54].

Collectively, these results establish AURA as the first framework capable of training ML models at unprecedented genomic scale, unlocking the potential for population-level resistance profiling and precision-guided treatment strategies in tuberculosis.

## Discussion

In this study, we introduce AURA, the first GPU-accelerated, de novo ML framework specifically designed for high-throughput prediction of antibiotic resistance in MTB. Using WGS data from 12,185 clinical isolates, AURA leverages massive GPU parallelization to deliver unprecedented scalability in resistance prediction. The workflow features GPU-optimized sparse data structures for constructing compressed sparse row (CSR) representations of *k*-mer frequency matrices, an efficient GPU-parallelized *χ*^2^ statistical test for robust univariate feature selection, and systematic training of advanced ML models across multiple *k*-mer lengths and hyperparameter configurations.

Previous frameworks relied on CPU-parallelized implementations that, while effective for small datasets, were unable to scale to tens of thousands of genomes due to prohibitive computational demands. For example, MTB++ required over 2 TB of disk space, hundreds of gigabytes of RAM, and nearly a month of compute time on a 100-core CPU cluster to process moderate datasets [54]. In contrast, AURA achieved a 6.2× speedup in feature matrix generation and a 215-fold acceleration in feature selection while reducing maximum memory usage by more than 100-fold. Crucially, these performance gains enabled training on the largest MTB resistance dataset to date, an advance that was previously computationally infeasible. End-to-end model training for clinically important antibiotics like isoniazid was reduced from 35 to less than 12 minutes, a threefold speed increase, while maintaining predictive accuracy comparable to established tools [27, 65].

The CRyPTIC consortium has previously developed a machine-learning pipeline for antibiotic resistance prediction using the same CRyPTIC dataset [9]. Both CRyPTIC and AURA therefore operate on an identical underlying dataset of paired genomic and phenotypic measurements, but they differ in how they construct features, select them, and define the learning target. At the feature-construction stage, both approaches use genome-wide, alignment-free *k*-mer representations with rarity filtering, where very low-frequency *k*-mers (occurring ≤ 5 times in the dataset) are discarded to reduce sequencing-error noise; however, the CRyPTIC machine-learning method generates *k*-mers directly from reads and compresses them into lossless co-occurrence “patterns” using a CPU/HPC-oriented workflow, whereas AURA constructs an explicit sparse *k*-mer count matrix in CSR format and is engineered for GPU-efficient matrix construction and manipulation. For feature selection, CRyPTIC applies an F-test against quantitative log_2_(MIC) values prior to fitting an optimized XGBoost regressor, while AURA performs GPU-parallel univariate *χ*^2^ screening (after binarizing the CSR matrix) to enable scalable filtering of extremely large *k*-mer spaces and to support multiple downstream classifiers. These choices reflect different prediction objectives: CRyPTIC is formulated primarily as MIC prediction (including explicit handling of MIC censoring) and then derives binary resistance calls by thresholding predicted MICs, whereas AURA directly optimizes binary resistance classification using CRyPTIC-provided susceptible/resistant labels. Finally, CRyPTIC reports training on remote CPU-cluster resources, and reproducing the full pipeline is computationally demanding due to large-scale *k*-mer generation and pattern compression; moreover, the trained per-drug models are not distributed as readily reusable artifacts, so applying the approach typically requires resource-intensive retraining.

The ability to train ML models at the population scale not only improves predictive power but also enables the identification of resistance determinants that were undetectable in smaller datasets due to limited statistical power. Upon alignment of significant *k*-mers, AURA recovered 151 loci that overlapped the CRyPTIC GWAS loci results [61] and identified 59 novel genomic loci not previously reported. Among these, *PE PGRS57* (Rv3514) emerged as a compelling candidate, associated with resistance to five antibiotics, including ethambutol and moxifloxacin. The *PE PGRS* genes have been implicated in immune evasion and host–pathogen interactions [7, 35], and mutations in *PE PGRS57* were recently linked to resistance against a novel antibiotic class [53]. Other family members, such as *PE PGRS47* and *PE PGRS41*, suppress autophagosome formation in infected macrophages [13, 51, 76], suggesting a broader role in MTB adaptation under drug pressure.

Additional novel associations involved *katG* with resistance to ethionamide, kanamycin, and bedaquiline. While *katG*’s role in isoniazid activation is well established [63], these associations may indicate compensatory or collateral mechanisms in multidrug-resistant strains. Similarly, *pncA*, the main determinant of pyrazinamide resistance [30, 41], showed unexpected associations with aminoglycosides and NRDs. These findings underscore how scaling ML training to tens of thousands of genomes provides the statistical power to detect subtle but clinically meaningful resistance signals that smaller datasets could not resolve. These associations should be interpreted with caution, as they are unlikely to be causal and are more plausibly explained by correlated resistance phenotypes .

Collectively, our results demonstrate how GPU-accelerated ML frameworks like AURA fundamentally shift the landscape of microbial genomics. By overcoming computational barriers and allowing a comprehensive analysis of large-scale genomic datasets, AURA facilitates highly accurate resistance prediction and the discovery of novel genomic markers. This positions AURA as a powerful tool for precision-guided tuberculosis therapy and global surveillance of drug-resistant strains.

## Limitations

The AURA models have limitations primarily driven by data quality and labeling. The predictive performance depends on the quality of input sequencing data, the accuracy and consistency of AST, and the clinical ECOFFs used to dichotomize MICs. These factors are important for isolates near decision thresholds and for antibiotics with unimodal MIC distributions, where substantial overlap between susceptible and resistant populations reduces class separability and, consequently, discriminative resolution. By contrast, antibiotics with bimodal MIC distributions exhibit cleaner separation and improved discrimination. In addition, trained AURA models identify patterns present in the current dataset; novel resistance variants absent from these data will not be recognized, highlighting the need for continuous updating as new genomes and labels become available. Further, as with other computational approaches, AURA is not a standalone basis for clinical decision making, and confirmatory laboratory AST remains required.

AURAis constrained by the scope of the CRyPTIC dataset, which includes only lineages 1 through 4 out of 10 recognized lineages and results for 13 antibiotics, so extending the approach to additional lineages and drugs will require well-curated datasets that link sequencing with robust phenotypic AST for those compounds. For some antibiotics such as NRDs, the number of resistant isolates is low, which limits power and can reduce the stability of inferred associations and performance estimates. Separately, reported performance depends on the epidemiological cutoff used to binarize AST, so results may shift under alternative cutoffs.

It should be emphasized that the novel associations identified (Table 2) are statistical correlations and should not be interpreted as evidence of causality. A key limitation is that some apparent cross-resistance patterns (see Figure 6 in the supplements) rather than direct mechanistic links: isolates resistant to second-line drugs are frequently also resistant to first-line drugs due to shared treatment histories and co-selection, which can introduce confounding and inflate associations across multiple drugs. Accordingly, we present these loci as candidate signals that warrant cautious interpretation and follow-up validation through functional or mechanistic studies and/or carefully controlled analyses in datasets better suited to extract correlated resistance patterns.

Another important consideration is that AURA extracts *k*-mers from assembled contigs rather than directly from raw reads. Because genomes were assembled from Illumina short reads, variation in read quality and assembly contiguity can affect which *k*-mers are recovered and, consequently, model performance. Assembly yields a consensus of the dominant population, so low-frequency variants may be lost. Thus, resistance confined to minor subpopulations may not be represented in contigs, reducing sensitivity and potentially under-calling resistance in mixed infections. Extending AURA to operate on read-derived *k*-mers without assembly is an important direction to assess this impact and evaluate allele-frequency–aware representations. In addition, incorporating more diverse sequencing data types, such as long-read or hybrid (short+long) datasets, could improve assembly completeness and variant recovery, ultimately boosting predictive performance.

Additionally, short-read sequencing and short-read assemblies often cannot uniquely resolve the repetitive *PE PPE* gene families, so *k*-mer signals overlapping these loci may arise from assembly artifacts. Therefore, any *PE PPE* associations reported should be treated as provisional candidates that require orthogonal confirmation through targeted validation and independent evidence of the underlying variant(s) before mechanistic interpretation.

Lastly, AURA currently relies on univariate independence tests, which can overlook interactions among *k*-mers; while multivariate modeling could improve performance by capturing joint effects, it typically incurs polynomial computational complexity in the total number of candidate *k*-mers and scales quickly with the number of variables considered, so scalable multivariate selection/inference would be needed to realize these gains efficiently.

## Software and Data Availability

The trained models are integrated into a software package named AURA, which is available at https://github.com/M-Serajian/AURA. This package can be executed on CPU (default) and GPU platforms and is configured and trained on assembled contigs supplied in FASTA format. However, a modest refactoring of the input parser would enable direct processing of an unassembled left or right pair of FASTQ format. Isolates and their phenotypic resistance data are annotated with ERR accession numbers, as documented in the CRyPTIC available at https://ftp.ebi.ac.uk/pub/databases/cryptic/release_june2022/reuse/CRyPTIC_reuse_table_20240917.csv. The forked version of Gerbil is available at https://github.com/M-Serajian/gerbil-DataFrame.

## Supplementary File

## Appendix 1: Summary of Phenotypic Drug Susceptibility Distribution for 13 Antibiotics

**Table 4:**
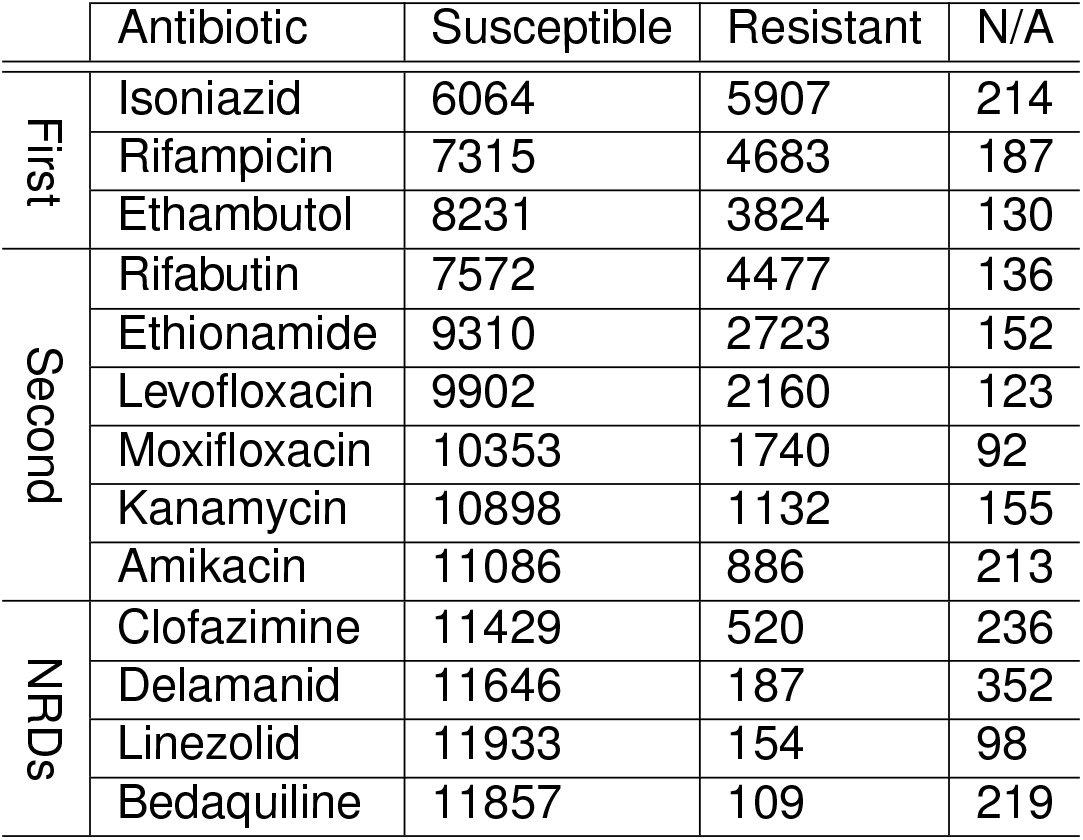
Distribution of susceptible, resistant, and unavailable phenotypes (N/A) isolates for each antibiotic in the CRyPTIC [61] dataset used for development of AURA.

## Appendix 2: Precision-Recall and Sensitivity-Specificity of AURA and Competing Methods

**Table 5:**
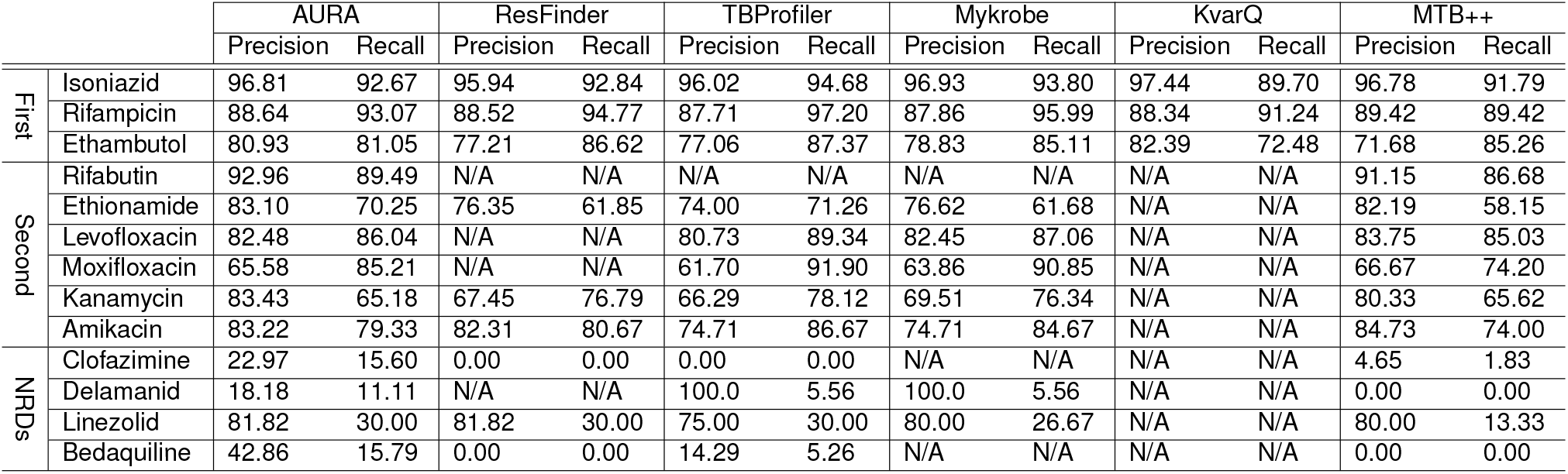
Precision and recall on the held-out test set for each antibiotic, evaluated with six resistance-prediction frameworks: AURA, ResFinder [19], TBProfiler [65], Mykrobe [27], KvarQ [58], and MTB++ [54].

**Table 6:**
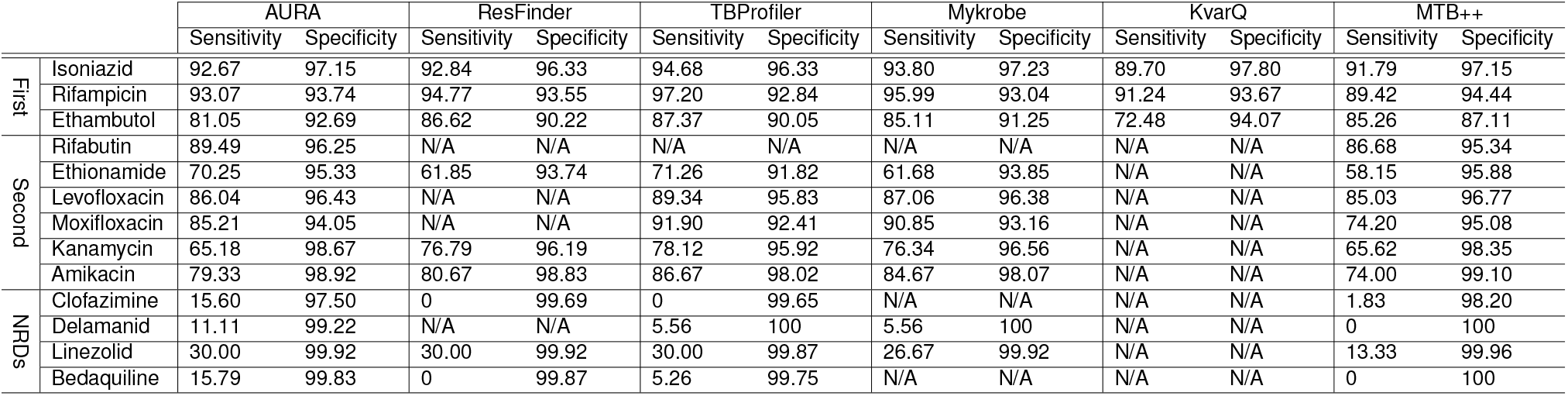
Sensitivity and specificity on the held-out test set for each antibiotic, evaluated with six resistance-prediction frameworks: AURA, ResFinder [19], TBProfiler [65], Mykrobe [27], KvarQ [58], and MTB++ [54].

## Appendix 3: F1-score Trajectories of AURA LR, RF, NB, XGBoost, SVM Across Various *k*-mer Length

**Figure 5:**
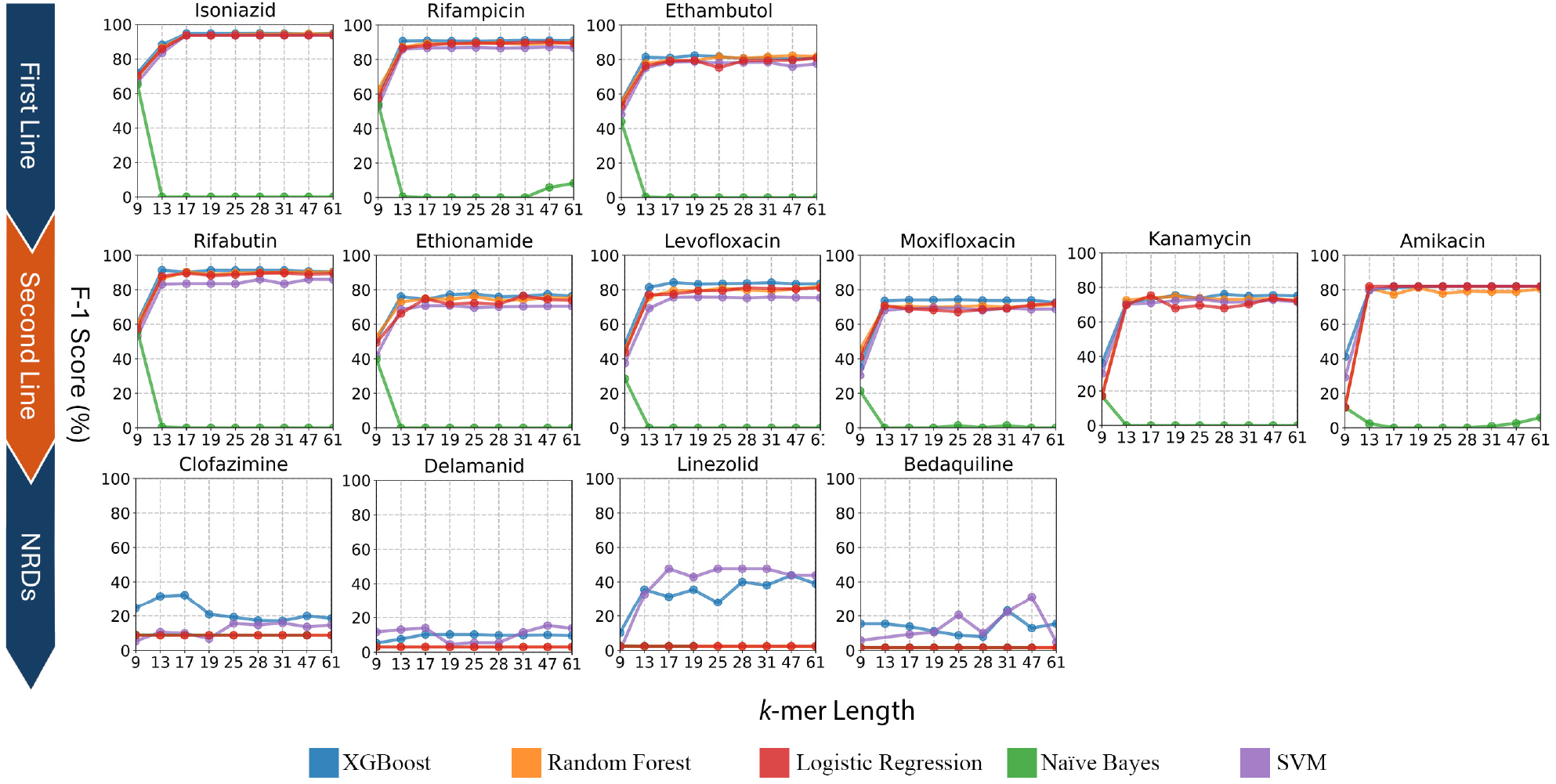
F1-score trajectories of AURA over six *k*-mer lengths (9, 13, 17, 19, 25, 28, 31, 47, 61 bp). For each antibiotic, the curve depicts the highest-performing classifier selected from XGBoost, SVM, RF, LASSO LR, or NB. The accompanying horizontal line, constant across all *k*-mer lengths, denotes the best competing method for the same drug. The results demonstrate the effect of *k*-mer length on the predictive performance of the models for first-line, second-line, and NRDs. For further details of the models and their parameters, see table 8 in the supplements.

## Appendix 4: Literature Review of the Associated Loci Found by AURA and Their Functionality in Other Studies

The *embB* gene is the most prevalent locus in this study, and the associations replicate identical patterns to CRyPTIC, linking *embB* to isoniazid, ethambutol, rifampicin, rifabutin, ethionamide, levofloxacin, and moxifloxacin. Ethambutol resistance arises when *embB* mutations alter the ara-binosyltransferase that builds the cell wall arabinan layer, thus reducing drug binding [57].

The ribosomal *rrs* gene is the next prevalent locus, matching CRyPTIC associations with ethionamide, levofloxacin, moxifloxacin, kanamycin, amikacin, and bedaquiline. The *rrs* gene encodes the 16S rRNA of the 30S ribosomal subunit—the binding site for several aminoglycosides, i.e. amikacin, kanamycin. The *rrs* mutations can modify the antibiotic-contact surface and undermine drug binding, a well-established mechanism of aminoglycoside resistance [1, 21, 28, 32, 61]. An interesting anlysis of the AURA reveal the single 31-mer aligned to *rrs* demonstrated a highly discriminative marker for amikacin resistance in this study: a linear model using this k-mer alone achieved an F1-score exceeding 80% on the held-out test set, underscoring its practical predictive value.

Likewise, the *gyrA* gene was revealed to be as the principal mechanism of resistance to most of the second-line antibiotics, echoing findings from the CRyPTIC consortium [61]. And the *gyrB* gene was exclusively identified for fluoroquinolone antibiotics, i.e. levofloxacin, moxifloxacin, identical to CRyPTIC. The *gyr* gene family in MTB encodes DNA gyrase, a type II topoisomerase essential for bacterial survival. DNA gyrase introduces negative supercoils into DNA in an ATP-dependent process and is required for replication, transcription, and recombination [31, 34, 38]. Because MTB lacks *parC* and *parE* homologs, DNA gyrase is the sole fluoroquinolone target in this organism [7, 38]. The enzyme consists of two gyrA and two gyrB subunits, encoded by *gyrA* and *gyrB*, respectively. Mutations in either gene can confer fluoroquinolone resistance, as shown in clinical and functional studies [6, 15, 24, 26]; however, such mutations appear far more frequently in *gyrA*. Yin et al. [77] reported that 73.3% of levofloxacin-resistant MTB isolates carried mutations in the *gyrA* QRDR, including 94.4% of strains with high-level resistance. Six single-codon substitutions (H70R, A90V, S91A, D94G, D94A, and D94N) and one double substitution (A90V + D94A) were reported. In contrast, only 1.6 % of isolates included *gyrB* mutation, T511N, in the absence of *gyrA* changes. These findings confirm that mutations at *gyrA* codons 90, 91, and 94 constitute the primary mechanism of fluoroquinolone resistance in MTB. It has been shown that the mutations in *gyrA* gene affects the enzyme’s binding pocket, sharply reducing fluoroquinolone affinity. As a result, fluoroquinolones, and often other second-line drugs, lose their effectiveness against MTB bacteria [33, 38, 61, 66, 68].

AURA also relied on *k*-mer aligned to the *fabG1* genetic region to predict resistance to six antibiotics, closely matching the CRyPTIC 2022 findings [61], except for clofazimine, which only CRyPTIC reported. The *fabG1* gene has been reported to be implicated in isoniazid resistance in MTB through promoter and coding-region mutations. The large-scale CRyPTIC 2024 study of more than 15,000 clinical isolates showed that the common *fabG1* c-15t promoter mutation markedly increases the minimum inhibitory concentration of isoniazid, often beyond epidemiological limits [9]. Resistance from this promoter change was comparable to, or greater than, that caused by *inhA* coding mutations (I21V, I21T). Other studies have also identified a synonymous coding-region change in *fabG1* (L203L) that creates an alternative promoter, drives *inhA* over-expression, and directly confers isoniazid resistance [2, 48, 64].

The *rpsL* gene was exclusively linked to resistance against all first-line antibiotics in our analysis, a pattern that matches the observations of the CRyPTIC study. The *rpsL* gene encodes the ribosomal protein S12, and mutations in this gene reshape the drug-binding site on the 30S ribosomal subunit. Streptomycin—one of the earliest anti-TB drugs—normally binds to this site and disrupts translation by inducing mRNA mis-reading. When the structure of S12 is altered, streptomycin can no longer bind effectively, so the antibiotic’s mechanism of action is dismantled and MTB survives under antibiotic pressure [12, 42].

## Appendix 5: Epidemiological Cut-off Values (ECOFF/ECVs) by CRyPTIC

**Table 7:** Proposed epidemiological cut-off values (ECOFF/ECVs) and suggested borderline MICs for selected compounds [10, 61] for the first-, second-line and NRDs.

| | Resistance Class | ECOFF/ECV $\text{mg L}^{-1}$ | Borderline MIC $\text{mg L}^{-1}$ |
| --- | --- | --- | --- |
| First | Isoniazid | 0.10 | 0.2, 0.4 |
|  | Rifampicin | 0.50 |  |
|  | Ethambutol | 4.00 | 4.00 |
| Second | Rifabutin | 0.12 |  |
|  | Ethionamide | 4.00 | 4.00 |
|  | Levofloxacin | 1.00 |  |
|  | Moxifloxacin | 1.00 |  |
|  | Kanamycin | 4.00 |  |
|  | Amikacin | 1.00 |  |
| NRDs | Clofazimine | 0.25 |  |
|  | Delamanid | 0.12 |  |
|  | Linezolid | 1.00 |  |
|  | Bedaquiline | 0.25 |  |

## Appendix 6: Optimized Models and Parameters for Each Antibiotic in AURA

**Table 8:** Optimized model configurations based on 70% training and 10% validation data. The table reports the selected model type, *k*-mer length, and number of features per antibiotic resistance class. These variables were optimized using the validation set after training on the training set. Final performance, shown in Tables 3 and 5, was evaluated on the 20% hold-out test set using these configurations for the first-line-, second-line, and NRDs.

| | Resistance Class | Model | $k$-mer length | Number of Features |
| --- | --- | --- | --- | --- |
| First | Isoniazid | XGBoost | 31 | 524288 |
|  | Rifampicin | XGBoost | 28 | 262144 |
|  | Ethambutol | XGBoost | 17 | 524288 |
| Second | Rifabutin | XGBoost | 28 | 524288 |
|  | Ethionamide | Random Forest | 25 | 262144 |
|  | Levofloxacin | XGBoost | 17 | 524288 |
|  | Moxifloxacin | XGBoost | 19 | 131072 |
|  | Kanamycin | XGBoost | 17 | 262144 |
|  | Amikacin | Random Forest | 19 | 32768 |
| NRDs | Clofazimine | XGBoost | 61 | 131072 |
|  | Delamanid | SVM | 61 | 131072 |
|  | Linezolid | SVM | 47 | 1024 |
|  | Bedaquiline | XGBoost | 31 | 131072 |

## Appendix 7: Cross-Resistance Analysis of The CrYPTIC Dataset

**Figure 6:**
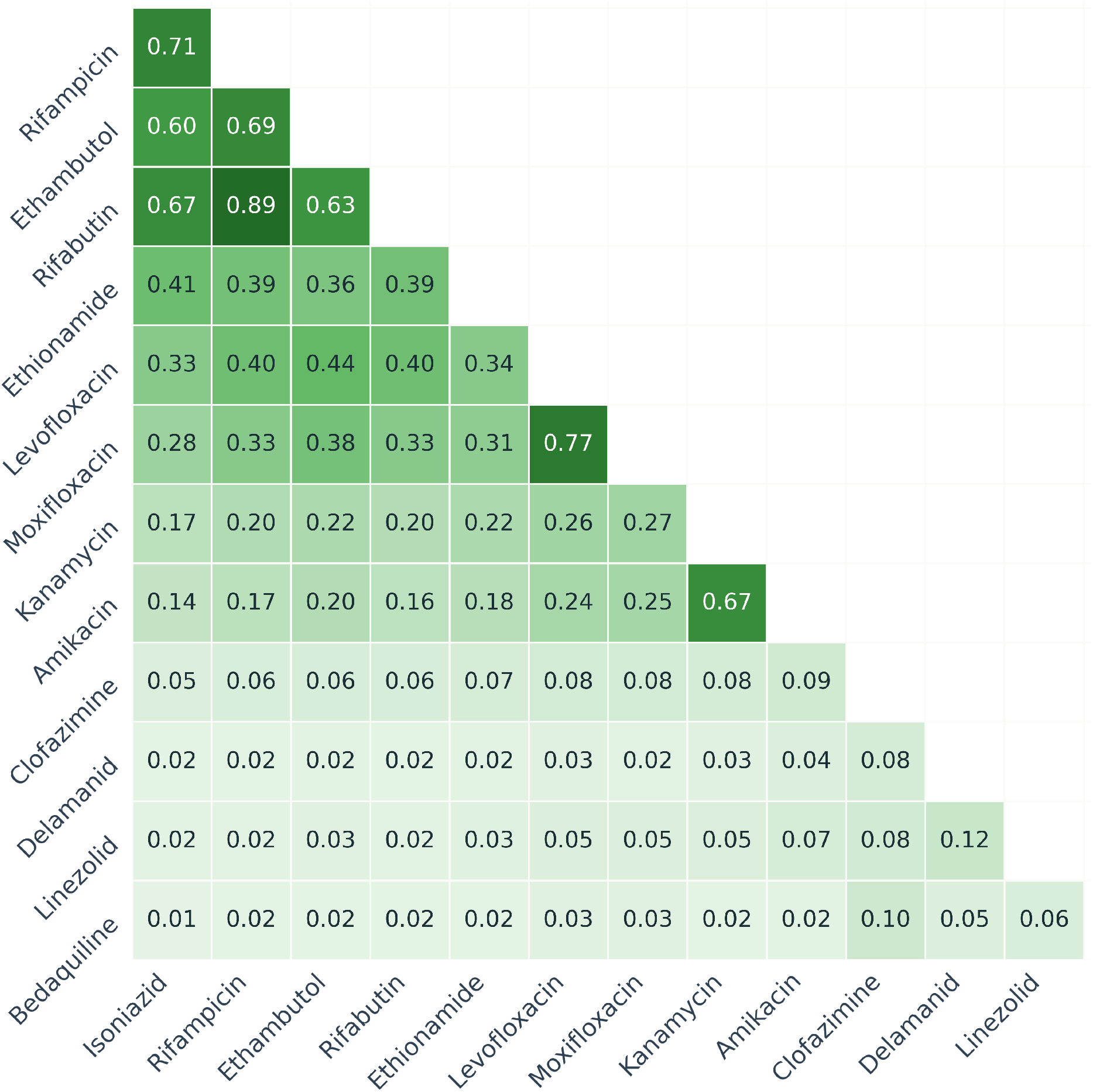
Cross-resistance patterns among antibiotic resistance phenotypes in the CRyPTIC dataset [8], quantified using the Jaccard similarity index. Each number represents the proportion of isolates resistant to both antibiotics relative to those resistant to either.

## Footnotes

1 CRyPTIC Consortium. *CRyPTIC reuse table (June 2022 release)*. European Bioinformatics Institute (EBI) FTP repository. Available at: https://ftp.ebi.ac.uk/pub/databases/cryptic/release_june2022/reuse/CRyPTIC_reuse_table_20240917.csv (accessed Jan 25, 2026).

## Notes

### Competing Interest Statement

The authors have declared no competing interest.

